# Delayed antibiotic exposure induces population collapse in enterococcal communities with drug-resistant subpopulations

**DOI:** 10.1101/766691

**Authors:** Kelsey M. Hallinen, Jason Karslake, Kevin B. Wood

**Affiliations:** Department of Biophysics, University of Michigan, Ann Arbor, USA; Department of Physics, University of Michigan, Ann Arbor, USA

## Abstract

Bacteria exploit a diverse set of defenses to survive exposure to antibiotics. While the molecular and genetic underpinnings of antibiotic resistance are increasingly understood, less is known about how these molecular events influence microbial dynamics on the population scale. In this work, we show that the dynamics of *E. faecalis* communities exposed to antibiotics can be surprisingly rich, revealing scenarios where–for example–increasing population size or delaying drug exposure can promote population collapse. Specifically, we combine experiments in computer-controlled bioreactors with simple mathematical models to reveal density-dependent feedback loops that couple population growth and antibiotic efficacy when communities include drug-resistant (β-lactamase producing) subpopulations. The resulting communities exhibit a wide range of behavior, including population survival, population collapse, or one of two qualitatively distinct bistable behaviors where survival is favored in either small or large populations. These dynamics reflect competing density-dependent effects of different subpopulations, with growth of drug-sensitive cells increasing but growth of drug-resistant cells decreasing effective drug inhibition. Guided by these results, we experimentally demonstrate how populations receiving immediate drug influx may sometimes thrive, while identical populations exposed to delayed drug influx (and lower average drug concentrations) collapse. These results illustrate that the spread of drug resistant determinants—even in a simplified single-species communities—may be governed by potentially counterintuitive dynamics driven by population-level interactions.

## INTRODUCTION

Antibiotic resistance is a growing public health threat (1). Decades of rapid progress fueled by advances in microbiology, genomics, and structural biology have led to a detailed but still growing understanding of the molecular mechanisms underlying resistance (2). At the same time, recent studies have shown that drug resistance can be a collective phenomenon driven by emergent community-level dynamics (3, 4). For example, drug degradation by a sub-population of enzyme-producing cells can lead to cooperative resistance that allows sensitive (non-producing) cells to survive at otherwise inhibitory drug concentrations (5, 6, 7). Additional examples of collective resistance include density-dependent drug efficacy (8, 9, 10, 11), indole-mediated altruism (12), and increased resistance in dense surface-associated biofilms (13). The growing evidence for collective resistance underscores the need to understand not just the molecular underpinnings of resistance, but also the ways in which these molecular-level events shape population dynamics at the level of the bacterial community.

A number of recent studies illustrate how a deeper understanding of microbial population dynamics can lead to improved strategies for stalling the emergence of resistance. These anti-resistance approaches exploit different features of the population dynamics, including competitive suppression between sensitive and resistance cells (14, 15), synergy with the immune system (16), precise timing of growth dynamics or dosing (17, 18), responses to subinhibitory drug doses (19), and band-pass response to periodic dosing (10). Resistance-stalling strategies may also exploit spatial heterogeneity (20, 21, 22, 23, 24, 25), epistasis between resistance mutations (26, 27), hospital-level dosing protocols (28, 29), and regimens of multiple drugs applied in sequence (28, 30, 18, 31, 19, 32) or combination (33, 34, 35, 36, 37, 38, 39, 40), which may allow one to leverage statistical correlations between resistance profiles for different drugs (41, 42, 43, 44, 39, 37, 45, 46, 47, 48). As a whole, these studies demonstrate the important role of community-level dynamics for understanding and predicting how bacteria respond and adapt to antibiotics. Despite the relatively mature understanding of resistance at the molecular level, however, the population dynamics of microbial communities in the presence of antibiotics are often poorly understood.

In this work, we investigate dynamics of *E. faecalis* populations exposed to (potentially time-dependent) in2ux of ampicillin, a commonly-used β-lactam. *E. faecalis* is a Gram-positive bacterial species found in the digestive tracts in many animals, including humans, yet *E. faecalis* is also an opportunistic pathogen that contributes to a number of clinical infections, including infective endocarditis, urinary tract infections, and blood stream infections (49, 50, 51, 52). β-lactams are among the most commonly used antibiotics for treating *E. faecalis* infections, though resistance is a growing problem (53). Resistance to ampicillin can arise in multiple ways, including by mutations to the targeted penicillin binding proteins or production of β-lactamase, an enzyme that hydrolyzes the β-lactam ring and renders the drug ineffective. While relatively rare, β-lactamase producing strains are associated with multi-drug resistant, high-risk enterococcal infections (54, 55, 56). In addition, because *E. faecalis* often produce low levels of the enzyme, β-lactamase producing strains may be undetectable by traditional laboratory tests (57), making them more widespread than originally believed. More generally, enzymatic drug degradation is widespread mechanism of antibiotic resistance and has been recently linked to cooperative resistance in *E. coli* (5) and *S. pneumoniae* (6). These enzyme-producing communities have the potential to exhibit rich and counterintuitive population dynamics, particularly when environmental conditions–such as drug concentration–are not constant.

In addition to the potential for enzyme-mediated resistance, *E. faecalis* populations exhibit density-dependent growth when exposed to a wide range of antibiotics (11). Increasing population density typically leads to decreased growth inhibition by antibiotics, consistent with the classical inoculum effect (IE) (8). However, β-lactams can also exhibit a surprising “reverse” inoculum effect (rIE) characterized by increased growth of the population at lower densities (11, 58). In *E. faecalis*, the reverse IE arises from density-dependent changes in local pH (11), which are associated with increased activity of ampicillin and related drugs (59). Similar growth-driven changes in pH (in antibiotic-free environments) have been shown to modulate intercellular interactions (60) and even promote ecological suicide in other microbial species (61). In addition to these in vitro studies, recent work shows that *E. faecalis* infections started from high- and low-dose inocula lead to different levels of immune response and colonization in a mouse model (62). If similar feedback loops between population density and growth exist in the presence of antibiotics, they could play a critical role in tipping populations towards survival or extinction and shape the fate of antibiotic-resistant communities.

In this work, we show that density-dependent feedback loops couple population growth and drug efficacy in *E. faecalis* communities comprised of drug-resistant and drug-sensitive cells exposed to time-dependent concentrations of antibiotic. By combining experiments in computer-controlled bioreactors with simple mathematical models, we demonstrate that these populations can survive, be driven to extinction, or exhibit bistability–where high-density populations survive and low-density populations collapse–depending on the initial population composition and the rate of drug influx. Surprisingly, mathematical models also predict—and experiments confirm—a region of “inverted bistability”, where low-density populations thrive while high-density populations collapse. Mathematically, these dynamics reflect competing environmental feedback loops between different subpopulations; increases in the size of the resistant subpopulation decrease the effective drug concentration, while increases in total population density potentiate the drug. Consistent with this picture, we show experimentally that decoupling total density from drug potentiation eliminates the “inverted bistability”. Finally, guided by predictions from the model, we experimentally show that there are certain scenarios where populations receiving immediate drug influx may eventually thrive, while identical populations exposed to delayed drug influx–which also experience lower average drug concentrations–are vulnerable to population collapse. These results illustrate that the spread of drug resistant determinants—even in a simplified single-species population—exhibits rich and counterintuitive dynamics.

## RESULTS

### Resistant and sensitive populations exhibit opposing density-dependent effects on antibiotic inhibition

To investigate the dynamics of *E. faecalis* populations exposed to β-lactams, we first engineered drug resistant *E. faecalis* strains that contain a multicopy plasmid that constitutively expresses β-lactamase (Methods). Sensitive cells harbored a similar plasmid that lacks the β-lactamase insert. To characterize the drug sensitive and drug resistant strains, we estimated the half maximal inhibitory concentration, IC_50_, of ampicillin in liquid cultures starting from a range of inoculum densities (Figure 1A; Methods). We found that the IC_50_ for sensitive strains is relatively insensitive to inoculum density over this range, while β-lactam producing resistant cells exhibit strong inoculum effects (IE) and show no inhibition for inoculum densities greater than 10^−5^ (OD units) even at the highest drug concentrations (10 µg/mL). To directly investigate growth dynamics at larger densities–similar to what can be resolved with standard optical density measurements–we used computer controlled bioreactors to measure per capita growth rates of populations held at constant densities and exposed to a fixed concentration of drug (as in (11)). At these higher densities, we found that resistant strains are insensitive to even very large drug concentrations (in excess of 10^3^ µg/mL). By contrast, sensitive populations are inhibited by concentrations smaller than 1 µg/mL, and the inhibition depends strongly on density, with higher density populations showing significantly decreased growth (Figure 1B)–indicative of a reverse inoculum effect (rIE). Taken together, these results illustrate opposing effects of cell density on drug efficacy in sensitive and resistant populations. In what follows, we focus on dynamics in the regime OD> 0.05, where the interplay between these two opposing effects may dictate survival or extinction of resistant populations.

**FIG 1.**
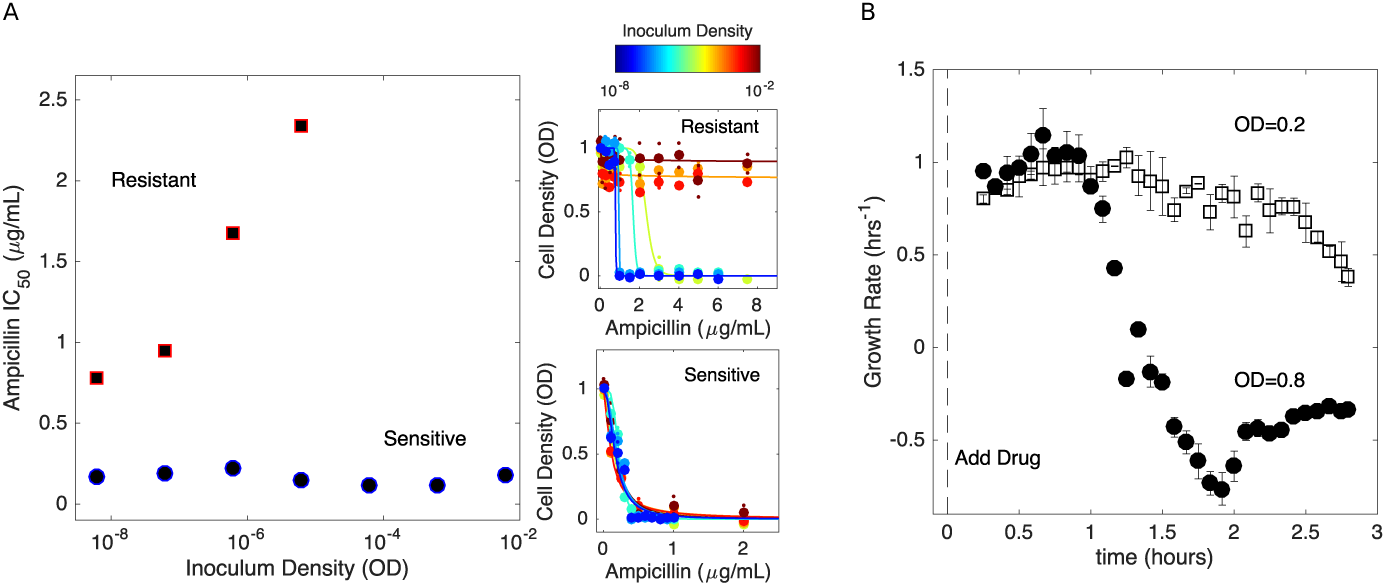
Changes in cell density have opposing effects on β-lactam eficacy in drug sensitive and drug resistant populations. A. Half-maximal inhibitory concentration (IC_50_) of ampicillin as a function of inoculum density for resistant (red squares) and sensitive (blue circles) populations. Right panels: dose response curves (cell density following 20 hrs of drug exposure vs. drug concentration) for drug resistant (top) and drug sensitive (bottom) populations at different inoculum densities (indicated by color, ranging from blue (low density) to red (high density)). Solid lines are fits to Hill-like function *f* (*x*) = (1 + (*x* /*K*)^*h*^)^−1^, where *h* is a Hill coefficient and *K* is the IC_50_. B. Per capita growth rate of drug sensitive populations held at a density of OD=0.2 (open squares) and OD=0.8 (filled circles) following addition of ampicillin at time 0. Growth rate is estimated, as in (11), from the average media flow rate required to maintain populations at the specified density in the presence of a constant drug concentration of 0.5 *µ*g/mL. Flow rate is averaged over sliding 20 minute windows after drug is added. Note that drug resistant populations exhibit no growth inhibition over these density ranges, even for drug concentrations in excess of 10^3^ *µ*g/mL.

### Resistant populations exhibit bistability between survival and extinction in the presence of constant drug influx

Bacteria in natural or clinical environments may often be exposed to drug concentrations that change over time. To introduce non-constant antibiotic concentrations, we used customized, computer controlled bioreactors capable of precise control of inflow (e.g. drug and media) and outflow in each growth chamber (Figure 2A; see also (63, 64, 11)). Cell density is monitored with light scattering (OD), and each chamber received fresh media and drug at a rate *µ* ≈ 0.12 hr^−1^, which is approximately an order of magnitude slower than the per capita growth rate of sensitive cells in drug-free media. In the absence of drug, cells reach a steady state population size of *C* (1 − *µ*), where *C* is the carrying capacity (*C* ≈ 1 in our experiments). By changing the concentration of drug *D*_*r*_ in the media reservoir, we can expose cells to effective rates of drug influx *F* = *µD*_*r*_.

**FIG 2.**
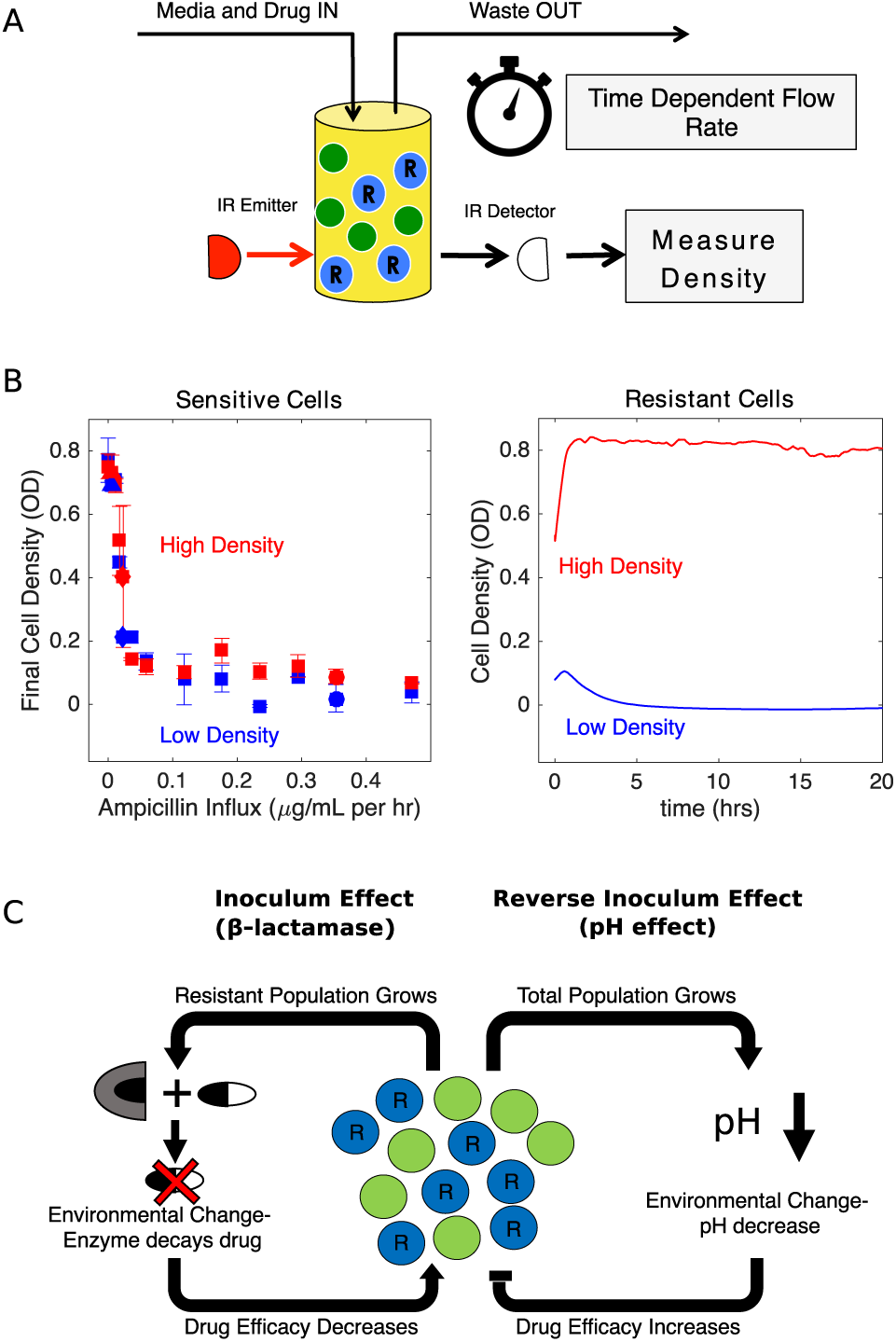
Homogeneous populations of drug resistant (but not drug sensitive) populations exhibit bistability between survival and extinction in the presence of constant β-lactam influx. A. Schematic of experimental setup. Cell density in planktonic populations is measured via light scattering from IR detector/emitter pairs calibrated to optical density (OD). Fresh media (containing appropriate drug concentrations) is introduced over time using computer-controlled peristaltic pumps, and waste is simultaneously removed to maintain constant volume (see Methods). B. Left panel: final cell density of drug sensitive populations exposed to constant drug influx over a 20 hour period. Experiments were started from either “high density” (OD=0.6, red) or “low density” (OD=0.1, blue) initial populations. Right panel: cell density time series for drug resistant populations exposed to ampicillin influx of approximately 1200 *µ*g/mL per hour. Experiments were started from either “high density” (OD=0.6, red) or “low density” (OD=0.1, blue) initial populations. In all experiments media was refreshed and waste removed at a rate of *µ* ≈ 0.12 hr^−1^, approximately 10 times slower than the per capita growth rate of cells in drug-free media. C. In mixed populations containing both sensitive (green) and resistant (blue, “R”) cells, there are opposing density-dependent effects on drug efficacy. Increasing the density of resistant cells is expected to decrease drug efficacy as a result of increased β-lactamase production (left side). By contrast, increasing the density of the total cell population decreases the local pH and increases the efficacy of β-lactam antibiotics (right side).

We first characterized the population dynamics of each cell type (resistant, sensitive) alone in response to different influx rates of ampicillin. In each experiment, we started one population at OD=0.6 (“high-density”) and one at OD=0.1 (“low density”). Not surprisingly, sensitive only populations exhibit a monotonic decrease in final (20 hour) population size with increasing drug concentration (Figure 2B, left panel), with both high and low-density populations approaching extinction at *D*_*r*_ < 1 µg/mL, corresponding to *F* < 0.1 µg/mL per hour. By contrast, high- and low-density populations of resistant cells exhibit strikingly divergent behavior, with high-density populations surviving and low-density populations collapsing (Figure 2B, right panel). In addition, we note that the resistant strains have dramatically increased minimum inhibitory concentrations (MIC), with high-density populations surviving at *D*_*r*_ = 10^4^ µg/mL (an effective influx of over 1000 µg/mL per hour). Indeed, the half-maximal inhibitory concentrations (IC_50_) for sensitive-only and resistant-only populations differ significantly even at very low densities (Figure 1), suggesting intrinsic differences in resistance even in the absence of density-dependent coupling. This difference corresponds to a direct benefit provided to the enzyme-producing cells, above and beyond any benefit that derives from drug degradation by neighboring cells.

These results, along with those in previous studies (11), are consistent with a picture of competing density-dependent feedback loops in populations comprised of both sensitive and resistant sub-populations (Figure 2C). Increasing the total population density potentiates the drug, a consequence of the pH-driven reverse inoculum effect (rIE). On the other hand, increasing the size of only the β-lactamase producing subpopulation is expected to decrease drug efficacy as enzymatic activity decreases the external drug concentration. These opposing effects couple the dynamics of different subpopulations with drug efficacy, which in turn modulates both the size and composition of the community.

### Mathematical model of competing density effects predicts bistability favoring survival of high-density populations at high drug influx rates and low-density populations at low influx rates

To investigate the potential impact of these competing density effects on population dynamics, we developed a simple mathematical model that ascribes density-dependent drug efficacy to a change in the effective concentration of the antibiotic, similar to (11). Specifically, the dynamics of sensitive and resistant populations are described by

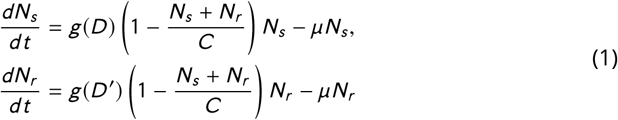

where *N*_*s*_ is the density of sensitive cells, *N*_*r*_ the density of resistant cells, *C* is the carrying capacity (set to 1 without loss of generality), *µ* is a rate constant that describes the removal of cells due to (slow) renewal of media and addition of drug, *D* is them effective concentration of drug (measured in units of MIC of the sensitive cells), and *D*′ = *D* /*K*_*r*_, where *K*_*r*_ is a factor that describes the increase in drug minimum inhibitory concentration (MIC) for the resistant (enzyme producing) cells in low density populations where cooperation is negligible. The function *g*(*x*) is a dose response function that describes the per capita growth rate of a population exposed to concentration *x* of antibiotic and is given by (9):

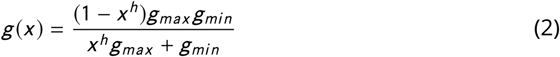

where *h* is a Hill coefficient that describes the steepness of the dose response function, *g*_*max*_ is the growth in the absence of drug (which we set to 1), and *g*_*min*_ > 0 is the maximum death rate. As drug concentration increases (*x* → ∞), the dose response function approaches this maximum death rate (*g* (*x*) → −*g*_*min*_).

To account for the density dependence of drug efficacy, we model the effective drug concentration as

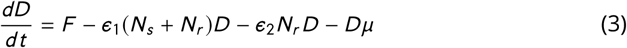

where *ϵ*_1_ < 0 is an effective rate constant describing the reverse inoculum effect (proportional to total population size) and *ϵ*_2_ > 0 describes the enzyme-driven “normal” inoculum effect (proportional to the size of the resistant subpopulation). *F* = *D*_*r*_ *µ* is rate of drug influx into the reservoir, which can be adjusted by changing the concentration *D*_*r*_ in the drug reservoir. When | *ϵ*_2_ | ≤ | *ϵ*_1_ | –when the per capita effect of the inoculum effect (IE) is less than or equal to that of its reverse (rIE) counterpart–the *ϵ*_1_ term is always larger in magnitude than the *ϵ*_2_ term and the net effect of increasing total cell density is to increase drug concentration, regardless of population composition. This regime is inconsistent with experiments, where resistant-only populations exhibit a strong IE and sensitive-only populations a rIE (Figures 1-2). We therefore focus on the case | *ϵ*_2_ | > | *ϵ*_1_ |, where density and composition-dependent trade-offs may lead to counterintuitive behavior.

Despite the simplicity of the model, it predicts surprisingly rich dynamics (Figure 3). At rates of drug influx below a critical threshold (*F* < *F*_*c*_), populations reach a stable fixed point at a density approaching *C* (1 − *µ*) as influx approaches zero. On the other hand, populations go extinct for large influx rates *F* ≫ *F*_*c*_, regardless of initial density or composition. Between the two regimes lies a region of bistability, where populations are expected to survive or die depending on the initial conditions. To characterize the behavior in this bistable region, we calculated the separatrix–the surface separating regions of phase space leading to survival from those leading to extinction–for different values of the antibiotic influx rate using an iterative bisection algorithm, similar to (65). The analysis reveals that increasing total population size can lead to qualitatively different behavior–survival or extinction–depending on the rate of drug influx.

**FIG 3.**
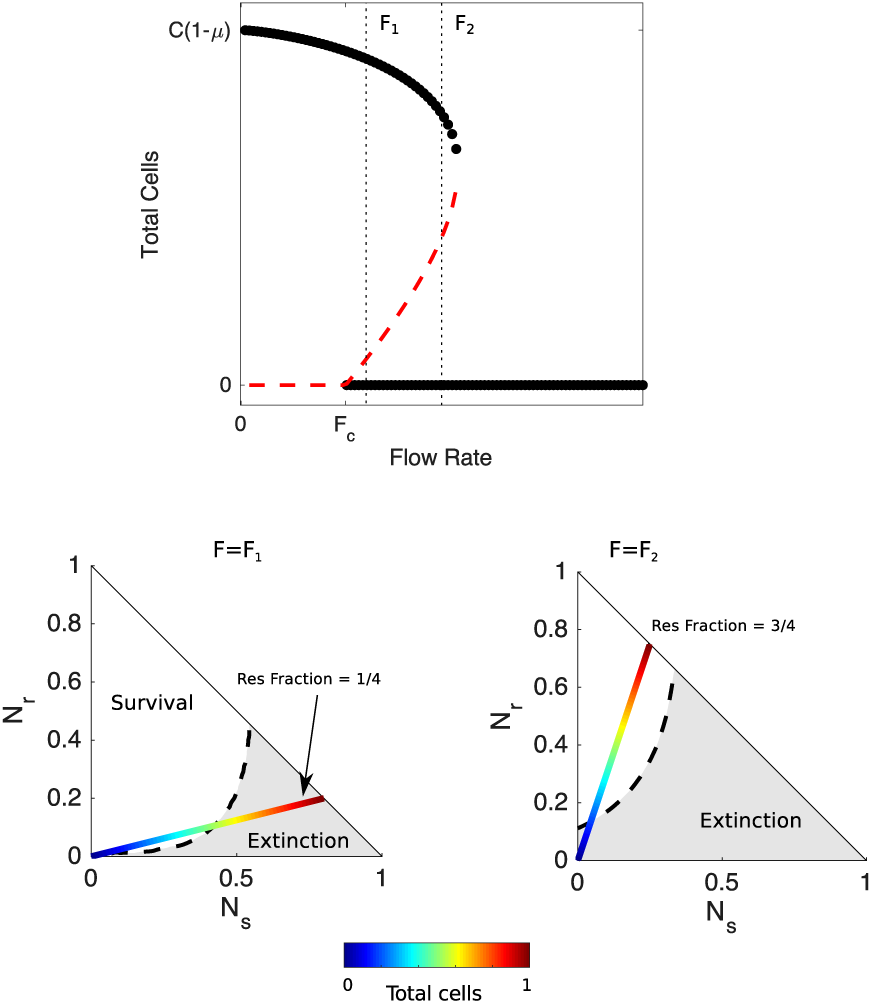
Mathematical model predicts bistability due to opposing density-dependent effects of sensitive and resistant cells on drug inhibition. Top: bifurcation diagram showing stable (filled circles) and unstable (red dashed curves) fixed points for different values of drug influx (*F*) and the total number of cells (*N*_*s*_ + *N*_*r*_). *F*_*c*_ ≈ *µK* is the critical value of drug influx above which the zero solution (extinction) becomes stable; *µ* is the rate at which cells and drugs are removed from the system, and *K*_*r*_ is the factor increase in drug MIC of the resistant strain relative to the wild-type strain. Vertical black dashed lines correspond to *F* = *F*_1_ > *F*_*c*_ (small drug influx, just above threshold) and *F*_2_ ≫ *F*_*c*_ (large drug influx). Bottom panels: regions of survival (white) and extinction (grey) in the space of sensitive (*N*_*s*_) and resistant (*N*_*r*_) cells for flow rate *F* = *F*_1_ (left) and *F* = *F*_2_ (right). Dashed lines show separatrix, the contour separating survival from extinction. Multicolor lines represent constant resistant fractions (1/4, left; 3/4, right) at different total population sizes (ranging from 0 (blue) to a maximum density of 1 (red)). Cell numbers are measured in units of carrying capacity. Specific numerical plots were calculated with *h* = 1.4, *g*_*min*_ = 1/3, *g*_*max*_ = 1, *є*_1_ = −1.1, *є*_2_ = 1.5, *γ* = 0.1, *K*_*r*_ = 14, *F*_1_ = 1.4, and *F*_2_ = 2.2.

For influx rates at the upper end of the bistable region–and for sufficiently high initial fractions of resistant cells–high-density populations survive while low-density populations go extinct (Figure 3, bottom right panel). For example, in populations with an initial resistant fraction of 3/4, small populations approach the extinction fixed point while large populations are expected to survive (Figure 3). Intuitively, the high-density populations have a sufficiently large number of resistant cells, and therefore produce a sufficient quantity of β-lactamase, that effective drug concentrations reach a steady state value below the MIC of the resistant cells, leading to a density-dependent transition from extinction to survival as the separatrix is crossed (Figure 3, bottom right).

Behavior in the low-influx regime of bistability (*F* ≈ *F*_*c*_) is more surprising. In this regime, the model predicts a region of bistability where initially high-density populations go extinct while low-density populations survive (Figure 3, bottom left). For example, at a resistant fraction of 1/4, low density populations will approach the survival fixed point while high density populations will approach extinction as the separatrix is crossed. These counterintuitive dynamics, which we refer to as “inverted bistability”, are governed in part by the reverse inoculum effect, which leads to a rapid increase in drug efficacy in the high-density populations and a corresponding population collapse. Mathematically, the different behavior corresponds to a translation in the separatrix curve as the influx rate is modulated (Figure 3; see also Figure S1).

To further characterize the dynamics of the model, we numerically solved the coupled equations (Equations 1, 3) for different initial compositions (resistant cell fraction) and different drug influx rates. In each case, we considered both high-density (OD=0.6) and low-density (OD=0.1) populations. As suggested by the bifurcation analysis (Figure 3), the model exhibits bistability over a range of drug influx rates (Figure 4A). The qualitative behavior within this bistable region can vary significantly. For small resistant fractions and low drug influx, bistability favors survival of low-density populations, while large resistant fractions and high drug influx favor survival of high-density populations. The parameter space is divided into 4 non-overlapping regions, leading to a phase diagram that predicts regions of extinction, survival, and bistabilities. It is notable that the dynamics leading to the fixed points can be significantly more complex than simple mononotic increases or decreases in population size (Figure 4A, top panels).

**FIG 4.**
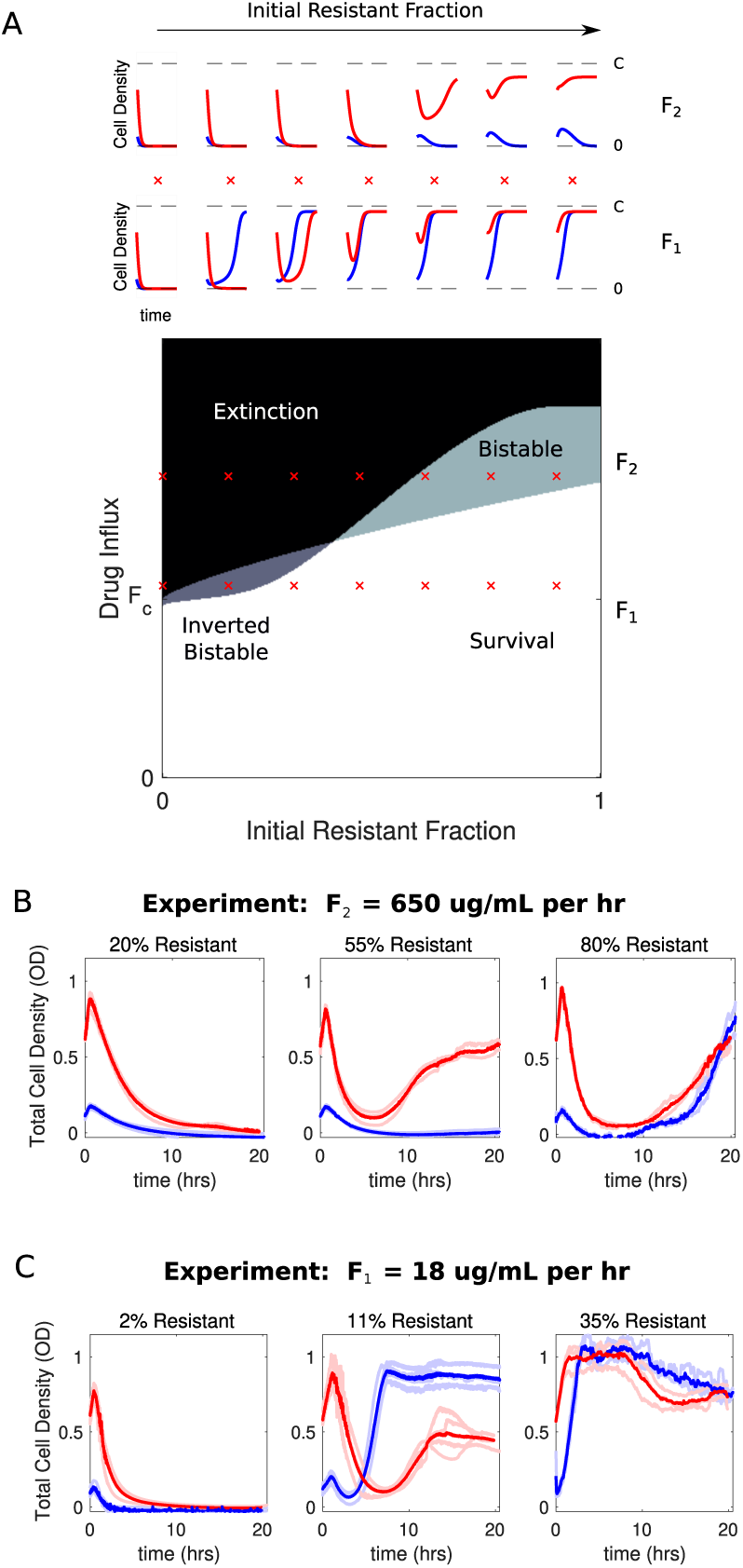
Bistability may favor survival of populations with highest or lowest initial density. A. Main panel: phase diagram indicating regions of extinction (black), survival (white), bistability (light gray; initially large population survives, small population dies), and “inverted” bistability (dark gray; initially small population survives, large population dies). Red ‘x’ marks correspond to the subplots in the top panels. Top panels: time-dependent population sizes starting from a small population (OD=0.1, blue) and large population (OD=0.6, red) at constant drug influx of *F*_2_ ≫ *F*_*c*_ (large drug influx) and *F*_1_ > *F*_*c*_ (small drug influx). *F*_*c*_ is the critical influx rate above which the extinct solution (population size 0) first becomes stable; it depends on model parameters, including media refresh rate (*µ*), maximum kill rate of the antibiotic (*g*_*min*_), the Hill coefficient of the dose response curve (*h*), and the MIC of the drug resistant population in the low density limit where cooperation is negligible (*K*). Specific numerical plots were calculated with *h* = 1.4, *g*_*min*_ = 1/3, *g*_*max*_ = 1, *ϵ*_1_ = −1.1, *ϵ*_2_ = 1.5, *γ* = 0.1, *K*_*r*_ = 14, *F*_1_ = 1.4, and *F*_2_ = 2.2. B. Experimental time series for mixed populations starting at a total density of OD=0.1 (blue) or OD=0.6 (red). The initial populations are comprised of resistant cells at a total population fraction of 0.2 (left), 0.55 (center), and 0.80 right) for influx rate *F*_1_ = 650 µg/mL. Light curves are individual experiments, dark curves are means across all experiments. C. Experimental time series for mixed populations starting at a total density of OD=0.1 (blue) or OD=0.6 (red). The initial populations are comprised of resistant cells at a total population fraction of 0.02 (left), 0.11 (center), and 0.35 right) for influx rate *F*_1_ = 18 µg/mL. Light curves are individual experiments, dark curves are means across all experiments.

### Small *E. faecalis* populations survive and large populations collapse when drug influx is slightly supercritical and resistant subpopulations are small

To test these predictions experimentally, we first performed a preliminary scan of parameter space in short, five-hour experiments starting from a wide range of initial population fractions and drug influx rates (Figure S2). Based on these experiments, we then narrowed our focus to a region of “high” influx rate (*F* ≈ 600 − 700 µg/mL per hour), where conditions may favor “normal” bistability, and a region of “low” influx rate (*F* ≈ 15 − 20 µg/mL per hour), where conditions may favor “inverted” bistability. Then, we performed replicate (*N* = 3) 20-hour experiments starting from a range of population compositions. Note that in the absence of density-mediated changes in drug concentration, these flow rates are expected to produce drug concentrations that increase over time, rapidly eclipsing the low-density limits for IC_50_’s of both susceptible and resistant cells (see Figure 1) and exponentially approaching steady state values of *D* = *F* /*µ* ≈ 8.5*F* with a time constant of *µ*^−1^ ≈ 8.5 hours.

The experiments confirm the existence of both predicted bistable regimes as well as the expected regimes of survival and extinction (Figure 4B-C). At each of the two flow rates (*F*_1_ and *F*_2_), we observe a transition from density-independent extinction–where populations starting from both high and low-densities collapse–to density-independent survival–where both populations survive–as the initial resistant population is increased (Figure 4B-C, left to right). However, in both cases there are intermediate regimes where initial population density determines whether the population will survive or collapse. When drug influx is relatively high (*F*_2_) and the population is primarily comprised of resistant cells (55 percent), initially large populations survive while small populations collapse (Figure 4 B, middle panel). On the other hand, when initial populations contain 11-15 percent resistant cells and drug influx is relatively small (*F*_1_), we observe a clear region of “inverted” bistability (Figure 4C, middle panel). In this regime, high-density populations (red) grow initially before undergoing dramatic collapse, while low-density populations (blue) initially decay before recovering and eventually plateauing near the carrying capacity.

### Inverted bistability depends on pH-dependent reverse inoculum effect

The model predicts that the inverted bistability relies on the reverse inoculum effect–Specifically, it requires ε_1_ < 0 and is eliminated when ε_1_ = 0 (Figure 5). Previous work showed that in this system, the reverse inoculum effect is driven by density-modulated changes in the local pH (11). Conveniently, then, it is possible–in principle–to eliminate the effect by strengthening the buffering capacity of the media. To test this prediction, we repeated the experiments in the inverted bistable region in strongly-buffered media (Figure 5). As predicted by the model, we no longer observe collapse of high-density populations, indicating that the region of inverted bistability is now a region of density-independent survival.

**FIG 5.**
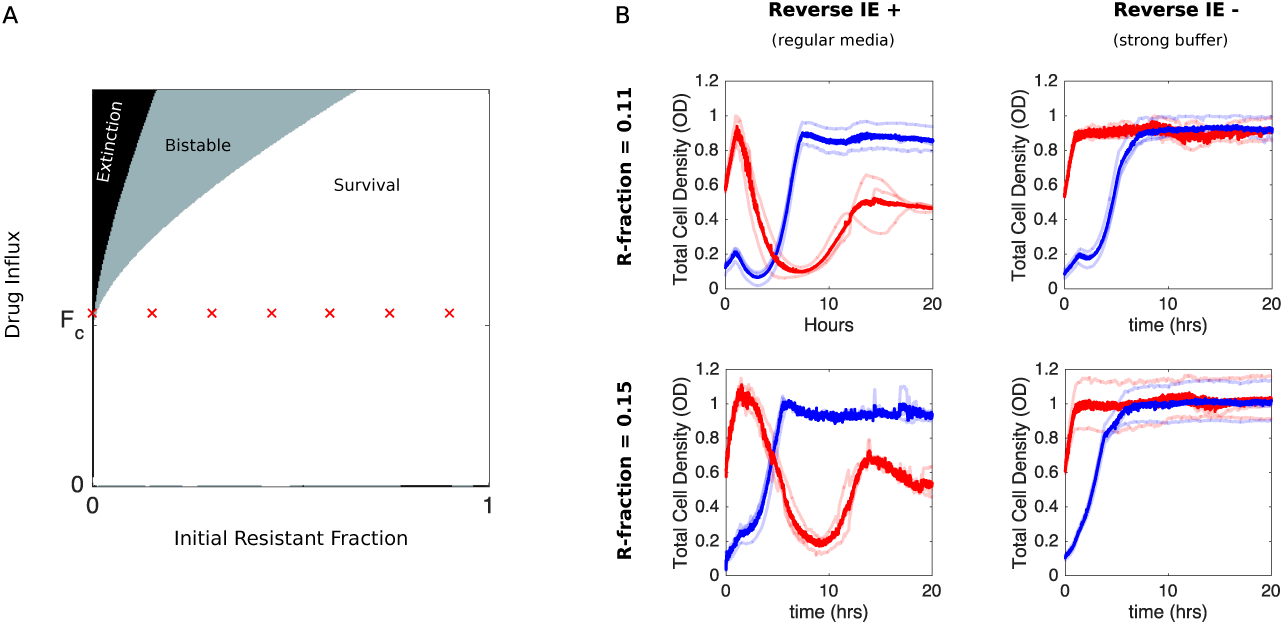
Eliminating reverse inoculum effect eliminates inverted bistability. A. Numerical phase diagram in absence of reverse inoculum effect (*ε*_1_ = 0) indicating regions of extinction (black), survival (white), and bistability (light gray; initially large population survives, small population dies). There are no regions of “inverted” bistability (initially small population survives, large population dies). Red ‘x’ marks fall along a line that previously traversed a region of inverted bistability in the presence of a reverse inoculum effect (Figure 4) but includes only surviving populations in its absence. *F*_*c*_ is the critical influx rate above which the extinct solution (population size 0) first becomes stable; it depends on model parameters, including media refresh rate (*µ*), maximum kill rate of the antibiotic (*g*_*min*_), the Hill coefficient of the dose response curve (*h*), and the MIC of the drug resistant population in the low density limit where cooperation is negligible (*K*). Specific phase diagram was calculated with same parameters as in Figure 4 except *e*_1_, which corresponds to the reverse inoculum effect, is set to 0. B. Experimental time series for mixed populations starting at a total density of OD=0.1 (blue) or OD=0.6 (red) in regular media (left panels) or strongly buffered media (right panels). The initial populations are comprised of resistant cells at a total population fraction of 0.11 (top) and 0.15 (bottom) and for influx rate of *F*_1_ = 18 µg/mL. Light curves are individual experiments, dark curves are means across all experiments.

### Delaying antibiotic exposure can promote population collapse

The competing density-dependent effects on drug efficacy raise the question of whether different time-dependent drug dosing strategies might be favorable for populations with different starting compositions. In particular, we wanted to investigate the effect of delaying the start of antibiotic influx for different population compositions and influx rates. Based on the results of the model, we hypothesized that there would be two possible regimes where delay could dramatically impact survival dynamics: one (corresponding to “normal bistability”) where delaying treatment would lead to larger end-point populations, and a second (corresponding to”inverted bistabillity”) where delaying treatment could, counterintuitively, promote population collapse (see Figure S3).

To test this hypothesis, we measured the population dynamics in mixed populations starting from an initial OD of 0.1 at time zero. We then compared final population size for identical populations experiencing immediate or delayed drug influx, with delay ranging from 0.5-2.5 hours. In experiments with non-zero delays, antibiotic influx was replaced by influx of drug-free media (at the same flow rate) during the delay period. In the first case, we chose a relatively small initial resistant fraction (0.11) and a relatively slow drug influx rate (*F* = 18 µg/mL), while in the second case we chose a larger initial resistant fraction (0.55) and a faster drug influx (*F* = 650 µg/mL).

Remarkably, we found that delaying treatment can have opposing effects in the two scenarios (Figure 6). At high drug influx rates and largely resistant populations, immediate treatment leads to smallest final populations (Figure 6, right panels), consistent with model predictions of bistability. Intuitively, the delay allows the subpopulation of resistant cells to increase in size, eventually surpassing a critical density where the presence of enzyme is sufficient to counter the inhibitory effects of antibiotic. On the other hand, at lower influx rates and lower initial resistant fractions, we find that immediate treatment leads to initial inhibition followed by a phase of rapid growth as the population thrives; by contrast, delays in treatment allow the population to initially grow rapidly before collapsing (Figure 6, left panels). It is particularly striking that delayed treatments–which also use significantly less total drug–can promote population collapse when immediate treatments appear to fail. Similar to the “inverted bistability” observed earlier, the beneficial effects of delayed treatment can be traced to density dependent drug efficacy–in words, the delay means the drug is applied when the population is sufficiently large that pH-mediated drug potentiation promotes collapse.

**FIG 6.**
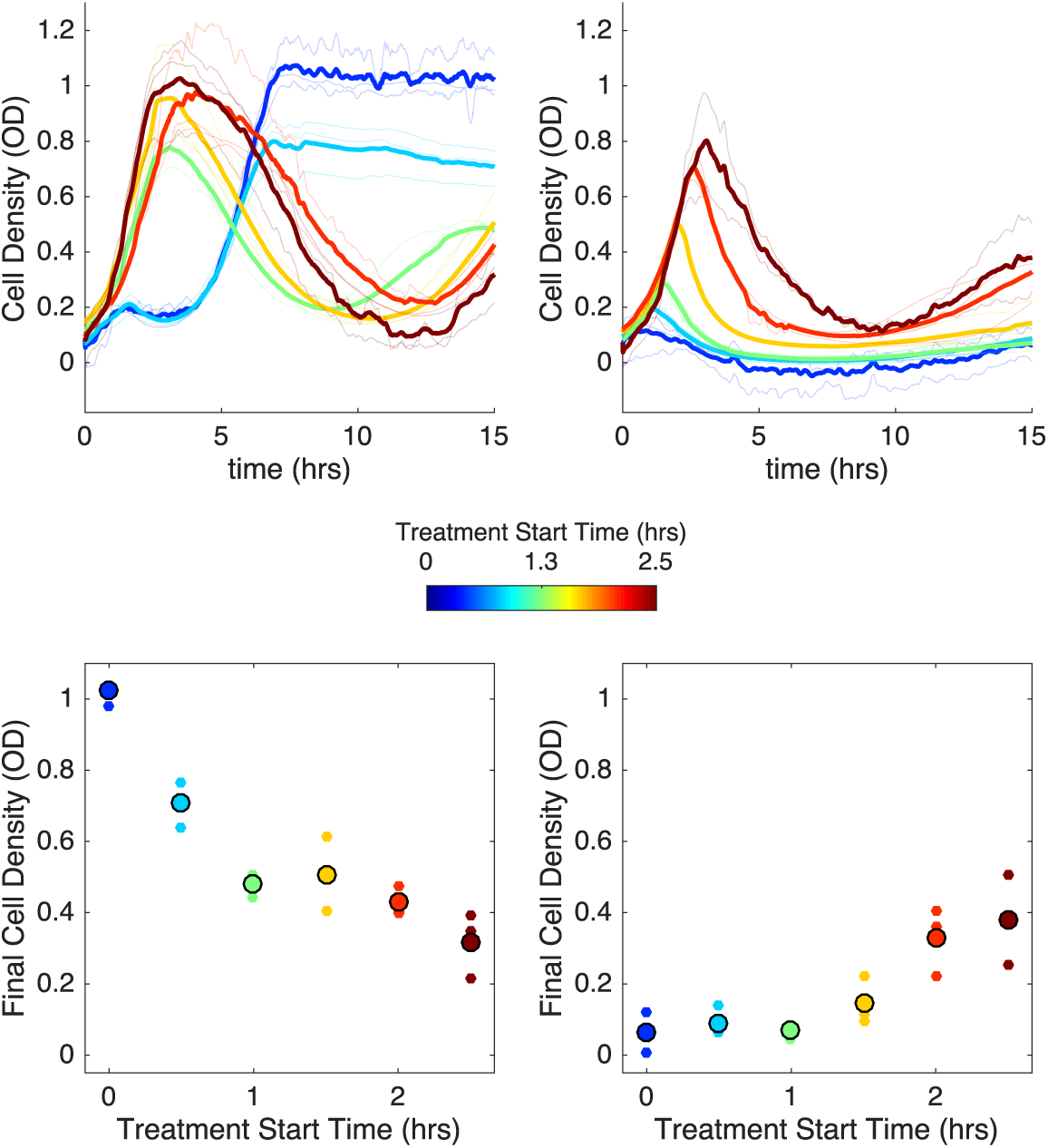
Delaying antibiotic exposure tips populations toward survival or extinction depending on initial resistance fraction and drug influx rate. Top panels: experimental time series for mixed populations with small initial resistance and low drug influx (left; initial resistance fraction, 0.11; *F* = 18 µg/mL) or large small initial resistance and high drug influx (right; initial resistance fraction, 0.55; *F* = 650 µg/mL). Antibiotic influx was started immediately (blue) or following a delay of up to 2.5 hours (dark red). Light transparent lines are individual replicates; dark lines are means over replicates. In experiments with nonzero delays, antibiotic influx was replaced by influx of drug-free media during the delay. Bottom panels: final cell density (15 hours) as a function of delay (“treatment start time”). small points are individual replicates; large circles are means across replicates.

## DISCUSSION

We have shown that different types of coupling between cell density and drug efficacy can lead to surprising dynamics in *E. faecalis* populations exposed to time-dependent ampicillin concentrations. In regimes of relatively fast or slow rates of drug influx, the results are intuitive: populations either survive or collapse, independent of initial population size (density). The intermediate regime, however, is characterized by bistability, meaning that population collapse will depend on initial population size. In regimes where cooperative resistance–in this case, due to enzymatic degradation of drug–dominates, larger populations are favored, similar to results predicted from the classical inoculum effect (9, 11). Under those conditions, it is critical to immediately expose cells to drug influx, as delays lead to increasingly resilient populations. Even more surprisingly, regimes characterized by comparatively smaller resistant populations and slower drug influx can lead to “inverted bistability” where initially small populations thrive while large populations collapse. In this case, delays to drug exposure can paradoxically promote population collapse. It is notable that the mathematical model suggests these results are not simply transient effects but instead reflect asymptotic behavior where the system approaches one of two stable fixed points (survival or extinction) with very different biological consequences.

Our goal was to understand population dynamics in simple, single-species populations where environmental conditions–including drug influx rate and population composition–can be well controlled. To make sense of experimental results and, more importantly, to generate new testable hypotheses, we developed a minimal mathematical model and analyzed its qualitative behavior using standard tools from dynamical systems and bifurcation theory. The model makes a number of surprising predictions about population survival and collapse which we were able to verify experimentally, suggesting that it captures some of the dominant behavioral features of the real system. However, we did not focus on quantitatively reproducing detailed features of the growth dynamics, which would likely require more detailed models with (potentially many) more parameters. For example, the model neglects evolutionary changes–for example, de novo mutations–that would certainly impact behavior on longer timescales. Similarly, we cannot definitely link all behavior in the experimental system to specific features of the model; it is possible, for example, that the observed population dynamics arise, at least in part, from mechanisms other than those considered here. For example, previous work (64) has shown that lysis of resistant cells can effectively increase the concentration of drug-degrading enzyme, and accounting for this effect may be required to reproduce biological dynamics under some conditions. Nevertheless, these results underscore the need for addition metrics (similar to the proposed notion of drug resilience (66)) that go beyond short-term growth measurements to for population dynamics over multiple timescales.

From an experimental perspective, it is obvious that the specific in vitro conditions used here fail to capture numerous complexities associated with resistance in clinical settings (67), including substantial spatial heterogeneity, potential for biofilm formation, effects of the host immune system, and drug concentrations that differ in both magnitude and time-course from the specific scenarios considered here. In addition, our experimental model system is based on plasmid-mediated resistance, and while this fact is not explicitly assumed in any of our mathematical models, horizontal gene transfer may introduce new dynamics (68, 69), particularly in high-density populations where conjugation is frequent. Nevertheless, the fact that population dynamics can be surprisingly complex, even under these simplified conditions, indicates that the spread of resistance alleles may not always follow simple selection dynamics. We therefore hope these results will motivate continued efforts to understand the potentially surprising ways that molecular level resistance events influence dynamics on the scale of microbial populations.

## METHODS

### Bacterial Strains, Media, and Growth Conditions

Experiments were performed with *E. faecalis* strain OG1RF, a fully sequenced oral isolate. Ampicillin resistant strains were engineered by transforming (70) OG1RF with a modified version of the multicopy plasmid pBSU101, which was originally developed as a fluorescent reporter for Gram-positive bacteria (71). The plasmid was chosen because it can be conveniently manipulated and propagated in multiple species (including *E. coli*) and contains a fluorescent reporter that provides a redundant control for readily identifying the strains. The modified plasmid, named pBSU101-BFP-BL, expresses BFP (rather than GFP in the original plasmid) and also constitutively expresses β-lactamase driven by a native promoter isolated from the chromosome of clinical strain CH19 (72, 73). The β-lactamase gene and reporter are similar to those found in other isolates of enterococci and streptococci (74, 75). Similarly, sensitive strains were transformed with a similar plasmid, pBSU101-DasherGFP, a pBSU101 derivative that lacks the β-lactamase insert and where eGFP is replaced by a brighter synthetic GFP (Dasher-GFP; ATUM ProteinPaintbox, https://www.atum.bio/). The plasmids also express a streptomycin resistance gene, and all media was therefore supplemented with streptomycin.

### Antibiotics

Antibiotics used in this study included Spectinomycin Sulfate (MP Biomedicals) and Ampicillin Sodium Salt (Fisher).

### Estimating IC_50_ for sensitive and resistant strains

Experiments to estimate the half-maximal inhibitory concentration (IC_50_) for each population were performed in 96-well plates using an Enspire Multimodal Plate Reader. Overnight cultures were diluted 10^2^ − 10^8^ fold into individual wells containing fresh BHI and a gradient of 6-14 drug concentrations. After 20 hours of growth the optical density at 600 nm (OD) was measured and used to create a dose response curve, which was fit to a Hill-like function *f* (*x*) = (1 + (*x* /*K*)^*h*^)^−1^ using nonlinear least squares fitting, where *K* is the half-maximal inhibitory concentration (IC_50_) and *h* is a Hill coefficient describing the steepness of the dose-response relationship.

### Continuous Culture Device

Experiments were performed in custom-built, computer-controlled continuous culture devices (CCD) as described in (11). Briefly, bacterial populations are grown in glass vials containing a fixed volume of 17 mL media. Cell density was measured at 1.5 second intervals in each vial using emitter/detector pairs of infrared LEDs (Radioshack). Detectors register a voltage output that is then converted to optical density using a calibration curve performed with a table top OD reader. Each vial contains input and output channels connected to silicone tubing and attached to a system of peristaltic pumps (Boxer 15000, Clark Solutions) that add drug and/or media and remove excess liquid on a schedule that can be programmed in advance or determined in real time. The entire system is controlled using a collection of DAQ and instrument control modules (Measurement Computing) and custom Matlab software (Mathworks) based on the Matlab Instrument Control Toolbox.

### Drug Dosing Protocols

In “constant flow” experiments, media (with drug, when relevant) is added at a rate of 1 mL/min for a total of 7.5 seconds every 3.75 minutes for an effective flow rate of 2 mL/hr (corresponding to a rate constant of *µ* = 0.12 hr^−1^ in 17 mL total volume). Media (plus cells and drug) is removed at an identical rate to maintain constant volume. While drug influx (and waste removal) strictly occurs on discrete on-off intervals, the timescale of those intervals (3-4 minutes) is an order of magnitude slower than the maximum bacterial growth rate under these conditions, which corresponds to a doubling time of approximately 30-40 minutes. The influx of drug is therefore approximately continuous on the timescale of bacterial dynamics. We experimentally modulate the influx rate of drug, *F*, without changing the background refresh rate (*µ*) by changing the drug concentration in the drug reservoir. For experiments involving time-dependent drug influx–for example, those in Figure 6, the media in the drug reservoirs is exchanged manually at specified times to mimic, for example, delayed treatment start times.

### Experimental Mixtures and Set up

All experiments were started from overnight cultures inoculated from single colonies grown on BHI agar plates with streptomycin and incubated in sterile BHI (Remel) with streptomycin (120 µg/mL) overnight at 37C. Highly buffered media was prepared by supplementing standard BHI with 50 µM Dibasic Sodium Phosphate (Fisher). Overnight cultures were diluted 100-200 fold with fresh BHI in continuous culture devices and populations were allowed to reach steady state exponential growth at the specified density (typically OD=0.1 or OD=0.6) prior to starting influx and outflow of media and waste. Experiments were typically performed in triplicate.

## ACKNOWLEDGEMENTS

This work is supported by the National Science Foundation (NSF No. 1553028 to KW; NSF GRF to KH), the National Institutes of Health (NIH No. 1R35GM124875-01 to KW), and the Hartwell Foundation for Biomedical Research (to KW). The format for this preprint is adapted from the American Society for Microbiology (ASM) template available on Overleaf.com.

## SUPPLEMENTAL MATERIAL

The Supplemental Material contains 3 supplemental figures (S1-S3).

**FIG S1.**
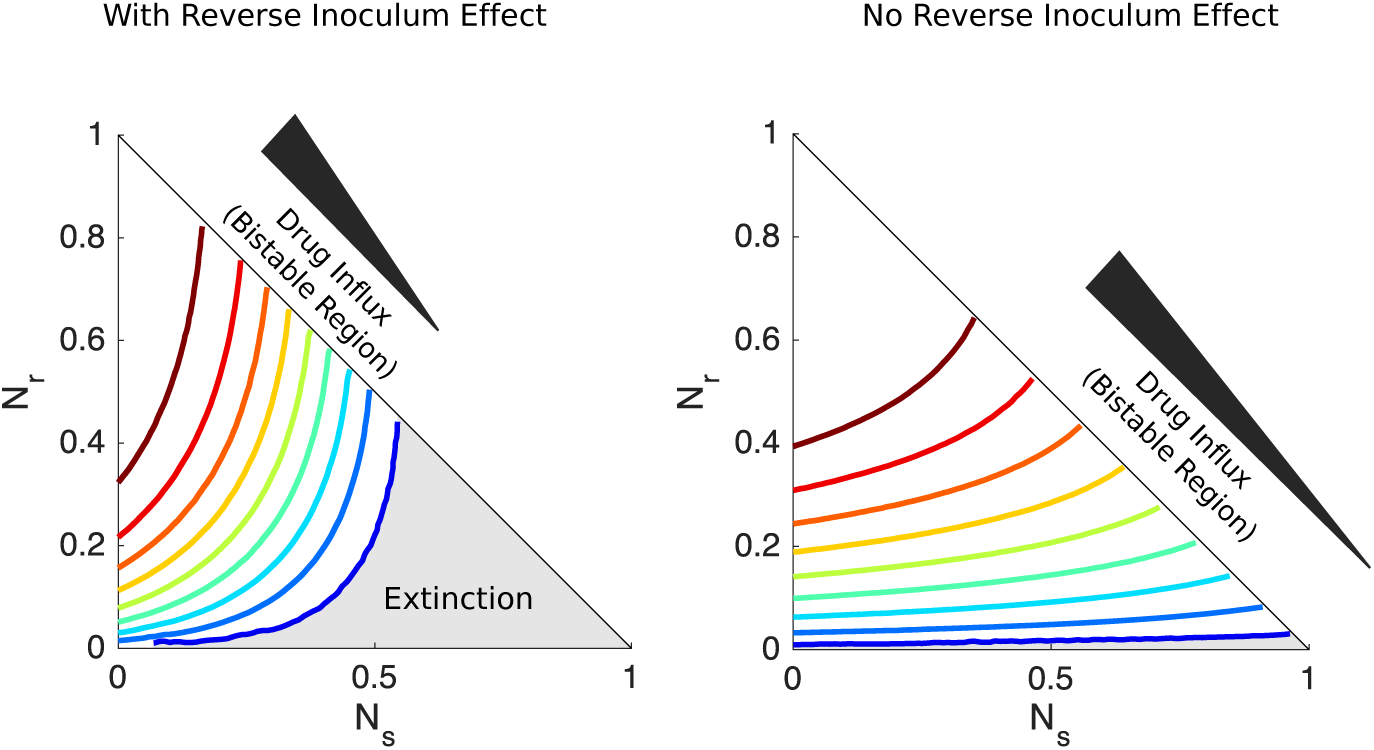
Separatrix contours separating survival from extinction with and without reverse inoculum effect. Contour lines separating regions of survival and extinction in the space of sensitive (*N*_*s*_) and resistant (*N*_*r*_) cells for a series of flow rates (increasing from blue to red) that span the bistable region with (left) and without (right) the reverse inoculum effect. Each contour line shows the separatrix for a particular value of *F* in the bistable region. For the lowest value of *F*, the region of extinction is shaded gray while the region of survival is white. Cell numbers are measured in units of carrying capacity. Specific numerical plots were calculated with *h* = 1.4, *g*_*min*_ = 1/3, *g*_*max*_ = 1, *ϵ*_2_ = 1.5, *γ* = 0.1, *K*_*r*_ = 14, and flow rates ranging from 1.4 < *F* < 2.7 (left) and 1.4 < *F* < 7.1 (right). *ϵ*_1_ = −1.1 in the left panel and 0 in the right panel. Initial drug concentration is set to 5x MIC of drug-sensitive cells in all panels, though qualitatively similar (but shifted) contours occur for other initial conditions.

**FIG S2.**
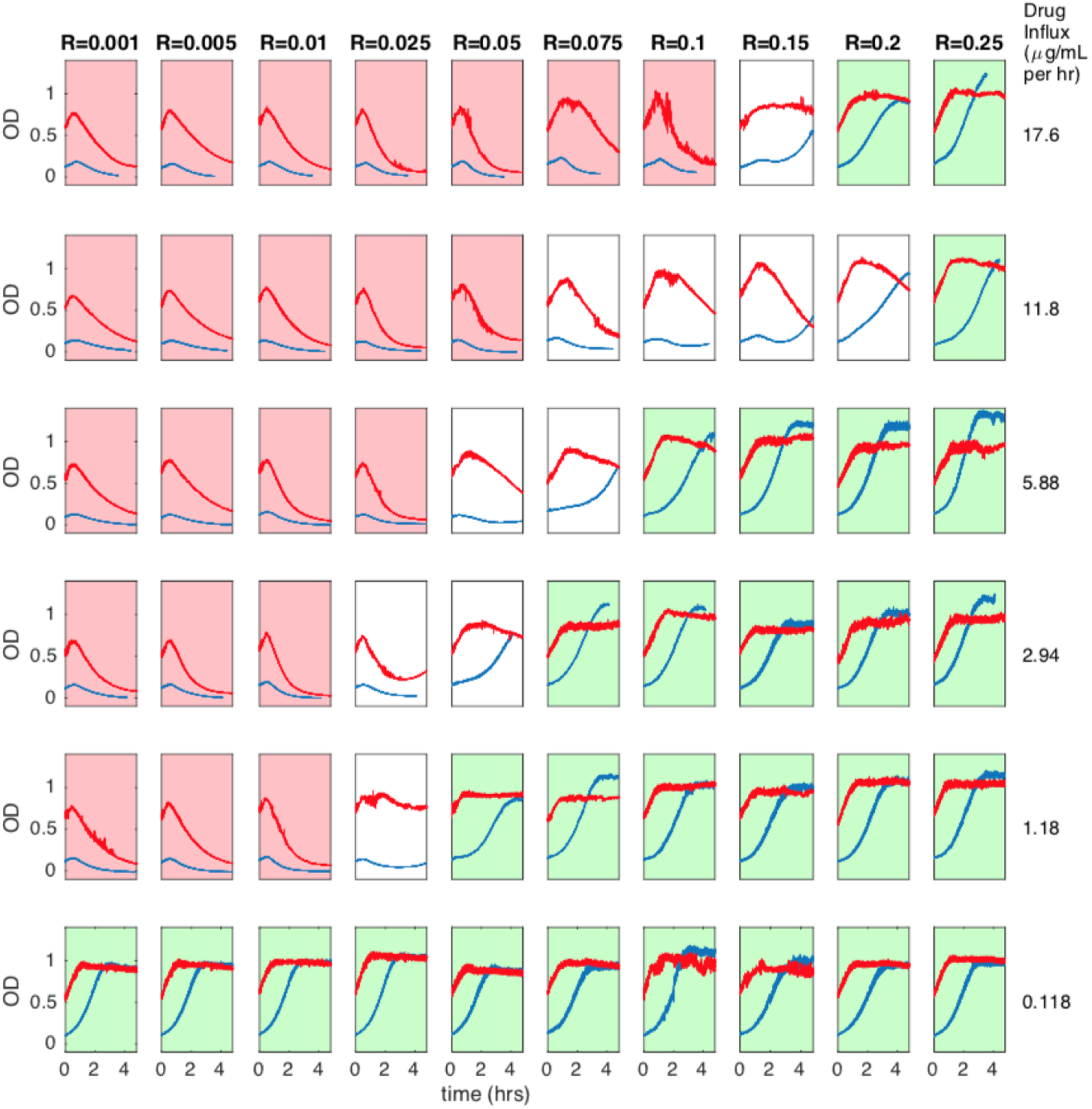
Short experiments to explore parameter space for inverted bistability. Cell density (OD) time series at various initial resistant fractions (columns) and ampicillin influx rate (rows). Populations were started at OD=0.1 (blue curves) or OD=0.6 (red curves). Red shaded plots indicate both populations are near extinction, while green shaded plots indicate that both populations are near carrying capacity at end of experiment (< 5 hrs).

**FIG S3.**
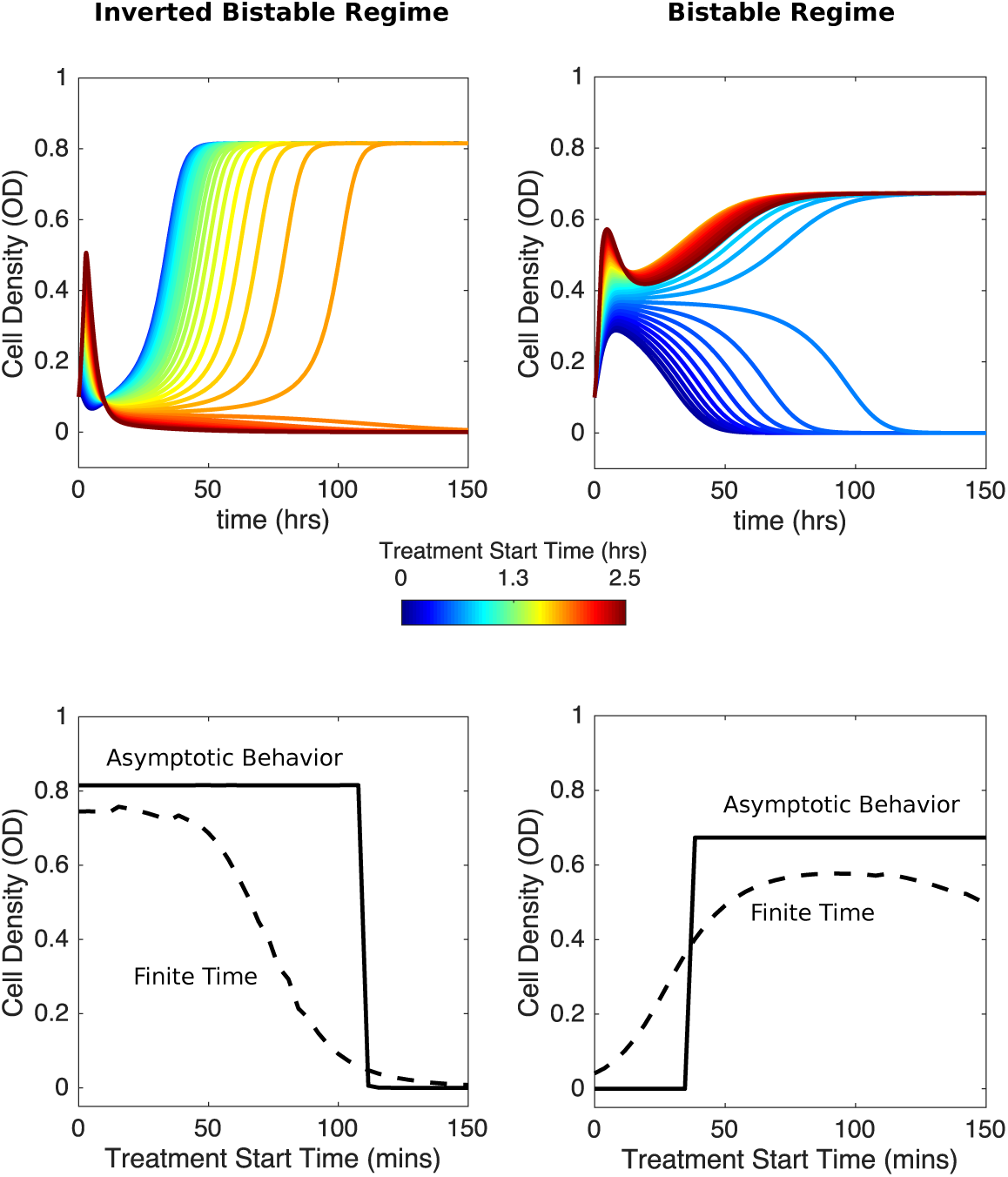
Numerical results inidcate that delaying antibiotic exposure tips populations toward survival or extinction depending on initial resistance fraction and drug influx rate. Top panels: time series for mixed populations with small initial resistance and low drug influx (left; initial resistance fraction, 0.04; *F* = 1.5) or large small initial resistance and high drug influx (right; initial resistance fraction, 0.56; *F* = 2.5). Antibiotic influx was started immediately (blue) or following a delay of up to 2.5 hours (dark red). Bottom panels: final cell density at finite times (40 hrs, dashed curves) or asymptotically long times (150 hrs; solid curves) as a function of delay (“treatment start time”). Model parameters are the same as those used in Figure 3.

